# A meta-analysis of Planarian: an animal model useful to understand the effect of magnetic field Planarian and magnetic field researches

**DOI:** 10.1101/252007

**Authors:** Liliana María Gomez Luna, Vivian de la Puente López

## Abstract

The planarians research since the first description until nowadays has a broad spectrum of topics. The present paper deals on a meta-analysis which identified as a problem the lack of consolidate information and data analysis about the use of planarian as animal model, and specifically the use of magnetic field in regeneration experiments. To conduct the meta-analysis 1255 papers published since 1955 to 2017 were analysed, identifying the Stem cell biology and regeneration as the most published topic, with 276 papers, followed by molecular and cellular analyses (238), genetics (175), inter and transdisciplinary research (107), ecotoxicological evaluations (102), animal model (67), ecological and biological studies (56), magnetic field (57), developmental biology (32) and RNA regulation (31).Other statistics and metrics indicators were taken in to account like total of papers and distribution per year, distribution of paper by journals and selection of main journals according to the number of published papers, most cited papers, authors and countries and distribution of papers by countries. Finally, were analysed those papers with Planarians research using magnetic field, all of them published during the last three years. It was an evidence that this topic is becoming a trending with rising interest, being the most reported species to study the magnetic field effects *Dugesia* spp. and *Girardia* sp.

**Summary statement:** This work presents a meta-analysis which allows consolidating information and a better understanding of trends in Planarian researches, emphasizing in the use of magnetic field.

## Introduction

Since 1970 with the advances in the information management and dissemination it is practically impossible to review all the scientific literatures even in an specific topic without the objectives and systematic strategies (Sánchez-Meca, 2010). Systematic reviews and, in particular, meta-analyses are a kind of scientific research aimed to objectively and systematically integrate the results of a set of empirical studies about a given research problem, with the purpose of determining the ¨state of the art¨ (Sánchez-Meca, 2010) in an specific research field. To accomplish that objective, carrying out a meta-analysis requires as the first step the formulation of the problem.

The present paper deals on a meta-analysis which identified as a problem the lack of consolidate information and data analysis about the use of planarian as animal model, and specifically the use of magnetic field in regeneration experiments in Planarian.

The birth of the study of planarians is most frequently associated with Pallas, who encountered them while exploring the Ural mountains in the late 18^th^ century, observing that these animals regenerate missing body parts after the fission (Pallas, 1774). However, other early reports exist, like the descriptions of Trembley, who described feeding pieces of planarians to hydra in his monograph in 1774 (Trembley, 1986) and Müller´s planarian species descriptions in 1773 (Müller, 1773).

Otherwise, Japanese encyclopaedias dating as far back as the 17^th^ century, describes land planarian (Elliott and Sánchez Alvarado, 2013), close to Geoplanidae family; but the oldest known reference to planarians comes from the Chinese text Yu-Yang Tsa-Tsu written around 860 AD by T’uan. He describes the animal “T’u-K’u” (likely the land planarian *Bipalium*), and hints at its regenerative abilities (Lue and Kawakatsu, 1986).

Planarians were described in the early 19^th^ century as being “immortal under the edge of the knife” and initial investigation of these remarkable animals was significantly influenced by studies of regeneration in other organisms and from the flourishing field of experimental embryology in the late 19^th^ and early 20^th^ centuries.

Having an almost unlimited capacity to regenerate tissues lost to age and injury, planarians have long fascinated naturalists many years ago (Elliott and Sánchez Alvarado, 2013). The planarians research since the first description until nowadays has a broad spectrum of topics. Some species were included in a number of experiments, all of them into the twelve genuses belonging to Dugesiidae family (Roskov Y., 2017).

The objective of this paper is to conduct a meta-analysis focused on Planarian researches to contribute with the understanding of the researches trends, but emphasizing in the use of magnetic field in regeneration process.

## Materials and methods

### Data sources and study selection criteria

The Sánchez-Meca methodology to meta-analysis was the theoretical referent to this research (Sánchez-Meca, 2010). To identify primary literature that examined the role of Planarian as animal model, main and trends topic and the place of research related with magnetic field effect a systematically search using Harzing´s Publish or Perish 5 software (Harzing, 2007) was done, with the following search terms: Planarian biological model* application*, Planarian magnetic field*, Planarian regeneration* Neoblast*, Planarian Ecology*, Planarian Toxicology*. The diagram with the meta-analysis strategy is presented in figure 1. We searched the references cited using Google scholar and specific search by database or by journal, reviewing the abstract when the whole paper was not available.

**Fig. 1.**
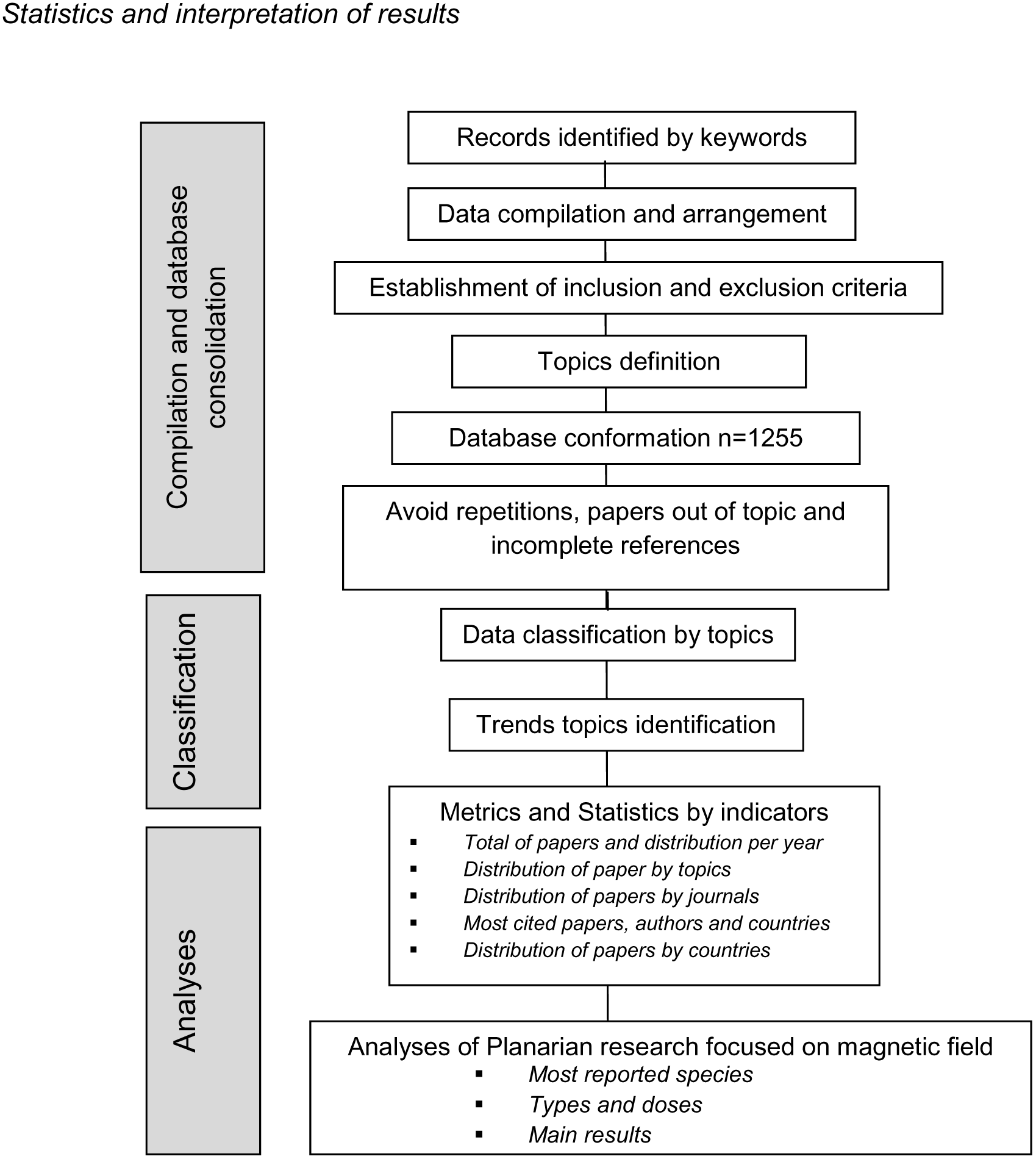
Diagram with the main steps of the study from compilation to the analyses.

### Study indicators

To analyse the database some indicators were considered:

– Topics (Main and trending topics)
– *Metrics and Statistics*
  ▪ Total of papers and distribution per year
  ▪ Distribution of paper by topics
  ▪ Distribution of papers by journals
  ▪ Most cited papers, authors and countries
  ▪ Distribution of papers by countries
– *Analyses of Planarian research focused on magnetic field*
  ▪ Most reported species
  ▪ Types and doses
  ▪ Main results

Statistics and interpretation of results

## Results and Discussion

### Total of papers and distribution per year

A Google scholar search using as a key word Planarian presents around 17.200 results in 0,52 s; if the key word is Planaria, there are less results: 16.200 in 0,11 s (94%); if Planarians 13.600 results in 0,07 s (79%). If the search is restricted to freshwater planarian 11.200 results appears in 0,08 s (65% in relation with Planarian results), while freshwater Planaria shows 9.050 results (0,07 s) (81%), but Planarian fundamentals only presents 1.400 results in 0,07 s (8%). Otherwise, while the use of Planarian worms as key word presents 11.100 results (0,09 s), Planarian flatworms shows 6.950 results (0,07 s) (63%) and surprisingly, Planarian flatworms regeneration 5.920 results (0,10 s), which represents the 85% of previous topic, while Planarian flatworm stem cell biology shows 2.390 results (0,14 s) (34%). It indicates the importance of Planarian regeneration as a research topic.

### Total of papers and distribution *per* year

Nevertheless, Planarian biology shows 21.400 results (0,11 s), even more than Planarian. If a specific search is developed about magnetic field and Planarians, 2.320 results (0,04 s) are showed, but if the key words are magnetic field and Planaria, only 1.930 (0,13 s) (83%), but surprisingly not all the papers are specific to Planarian as we present at the end of the analyses. It means that the search strategy needs to be well designed in order to have an adequate universe of results. In this case, according to the objective of this work, some descriptors or key words were selected, privileging the use of Planarian.

The search strategy using Publish or Perish included the following keywords: Planarian application, Planaria as biomodel, Planaria culture, Planarian and magnetic field, and Planaria and magnetic field, Planarians stem cell and neoblast regeneration, only in the title. With the entire data base consolidated the next step was to avoid the repetitions, removing the papers out of topic or with incomplete references.

As a result of the analysis it was conformed a database including 1255 papers. It was showed an increasing interest in Planarian researches since 1955 to 2017, being relevant the publishing shooting since 2010, to which correspond the 74% of the total papers reviewed (Fig. 2).

**Fig. 2.**
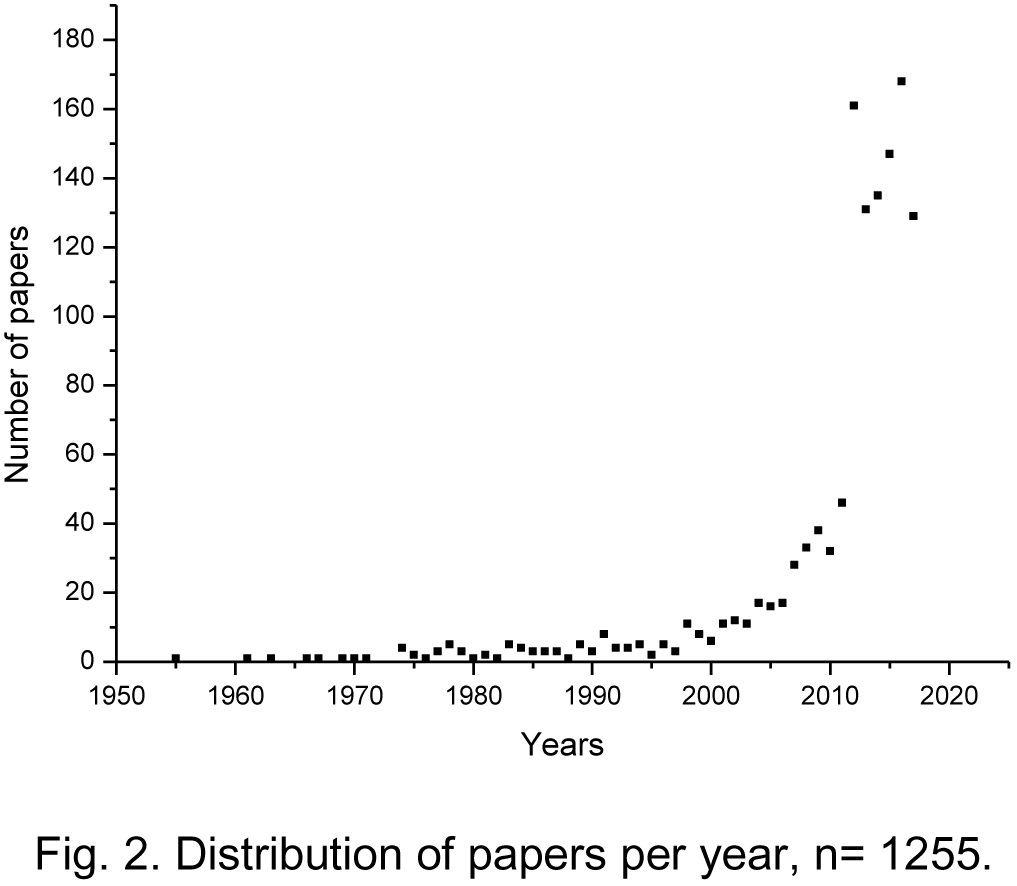
Distribution of papers per year, n= 1255.

According to a general analysis of all the papers included in this meta-analysis, it is possible to consider three stages in the planarian published papers: the first one includes papers from 1955 to 2000, the second from 2001 to 2010 and the last one from 2011 to 2017 (Table I).

**Table I.** Distribution of papers by topics and stages.

| Topics | Stage 1 | Stage 2 | Stage 3 | Total |
| --- | --- | --- | --- | --- |
| Stem cell biology and regeneration | 18 | 47 | 213 | 276 |
| Molecular and cellular analysis | 14 | 23 | 195 | 232 |
| Genetics | 10 | 37 | 128 | 175 |
| Inter and transdisciplinary research | 6 | 32 | 69 | 107 |
| Ecotoxicological evaluations | 14 | 13 | 75 | 102 |
| Animal model | 11 | 11 | 45 | 67 |
| Ecological and biological studies | 11 | 13 | 32 | 56 |
| Magnetic field | 6 | 3 | 48 | 57 |
| Development Biology | 2 | 10 | 20 | 32 |
| RNA regulation | 1 | 3 | 27 | 31 |
| Physiological evaluations | 6 | 9 | 11 | 26 |
| Morphology | 5 | 5 | 13 | 23 |
| Phylogenetic studies | 2 | 4 | 13 | 19 |
| Behaviour and planarian intelligence | 0 | 1 | 14 | 15 |
| Immunological studies | 2 | 3 | 10 | 15 |
| Bioinformatics | 1 | 1 | 10 | 12 |
| Proteomic analysis | 0 | 0 | 8 | 8 |
| Exobiology | 0 | 0 | 2 | 2 |
| **Total** | **109** | **215** | **933** | **1255** |
| **Percentage** | **8,7** | **16,9** | **74,3** |  |

### Distribution of paper by topics

There were identified 18 main topics. The growing interest since 2011 was not focused in a specific one, but all the topics, highlighting that the number of papers was highest in those related with stem cell biology and regeneration (213), molecular and cellular studies (195) and genetics (128) (Table I, Fig. 3). It is the same pattern of the total published papers.

**Table II.** Published papers by topics in the last 3 years (2015-2017).

| Topics | 2017 | 2016 | 2015 |
| --- | --- | --- | --- |
| Animal model in transdisciplinary researches | 5 | 7 | 6 |
| Immunological studies | 4 | 0 | 1 |
| Developmental biology | 1 | 4 | 3 |
| Behaviour and planarian intelligence | 2 | 2 | 6 |
| Bioinformatics | 3 | 1 | 4 |
| Ecological studies | 2 | 5 | 4 |
| Ecotoxicological evaluations | 8 | 20 | 10 |
| Exobiology | 1 | 0 | 0 |
| Genetics | 19 | 27 | 21 |
| Interdisciplinary research | 11 | 12 | 9 |
| Magnetic field | 8 | 7 | 6 |
| Molecular and cellular studies | 29 | 39 | 30 |
| Morphology | 3 | 1 | 3 |
| Phylogenetic studies | 1 | 2 | 4 |
| Physiological evaluations | 1 | 2 | 1 |
| Proteomic analyses | 1 | 1 | 1 |
| RNAs regulation | 7 | 2 | 1 |
| Stem cell biology and regeneration | 24 | 36 | 37 |

**Fig. 3.**
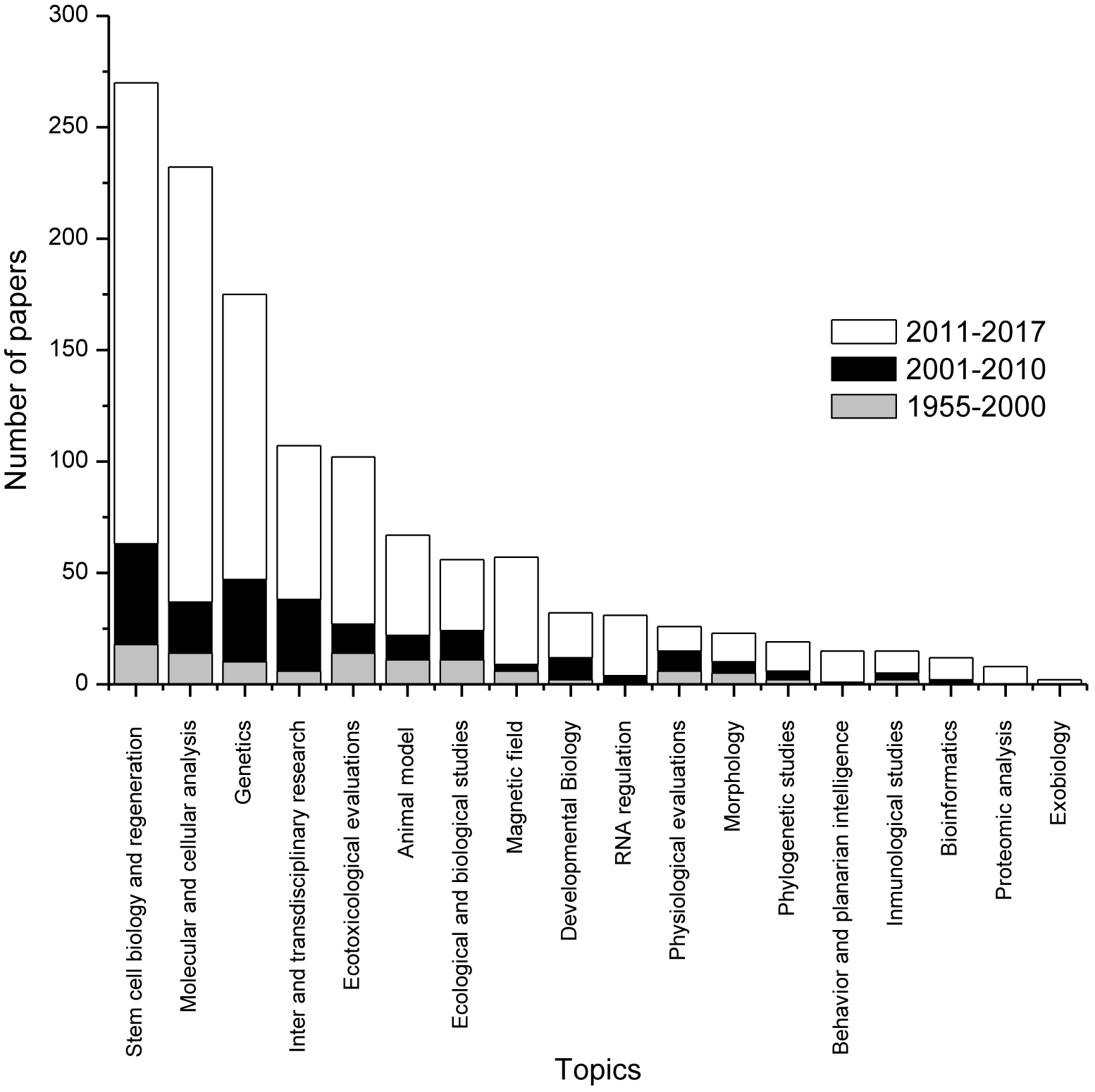
Variation of the number of publishing papers by topics into the three stages, n= 1255.

As a general comment the planarians researches have an arising interest since 1955 until nowadays (Fig. 2) and the specific analysis revealed the following main aspects:

– During the first two stages there was a gap in two topics: Proteomic analysis and Exobiology, these become really important just in the last stage (2011-2017), but have a few of published papers: 8 and 2, respectively.
– The topic related with the study of behaviour and planarian intelligence was growing from 0 to 15 published papers across the three stages.
– Practically there is a growing tendency of published papers to all the topics, which is coherent with the arising interest in Planarian research.
– The last stage is more diversify and new topics arise or have a strong growing tendency, like magnetic field researches, with 48 papers in the last stage and RNA regulation with 27.

A general analysis allows to identify the Stem cell biology and regeneration as the most published topic (276 papers), followed by molecular and cellular analyses (238), genetics (175), inter and transdisciplinary research (107), ecotoxicological evaluations (102), animal model (67), ecological and biological studies (56), magnetic field (57), developmental biology (32) and RNA regulation (31). This hierarchy is the trend in the last stage too, but researches about magnetic field arise to reach the sixth place.

### Analysis of trending topics

The trending topics were the topics identified during the last three years, which have a growing interest and a representative quantity of paper published. Considering the results presented in table II the following ten trending topics were selected:

1. Immunological studies
2. Bioinformatics
3. Ecotoxicological evaluations
4. Exobiology
5. Genetics
6. Interdisciplinary research
7. Magnetic field
8. Molecular and cellular studies
9. RNAs regulation
10. Stem cell biology and regeneration

### Distribution of papers by journals

In order to optimize time and enforcement this analysis considers a sample of 788 papers published in 360 journals, which included the 20 most cited papers.

The journals with more papers published are: Developmental Biology (30 papers), International Journal of Developmental Biology (29) and PLOS ONE 10^th^ Anniversary (27); following by Development Growth & Differentiation (18), PLOS Genetics (15), Developmental Cell (13) and Development Genes and Evolution (12) and Ecotoxicology and Environmental Safety (10). The rest of journals had 1 to 9 papers published (Table III).

**Table III.**
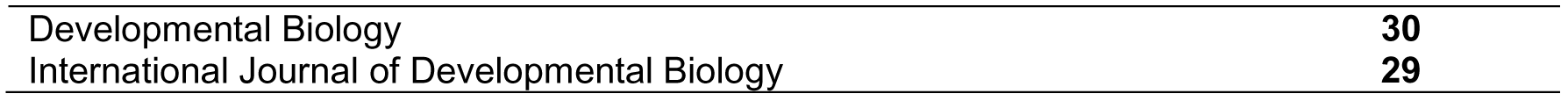
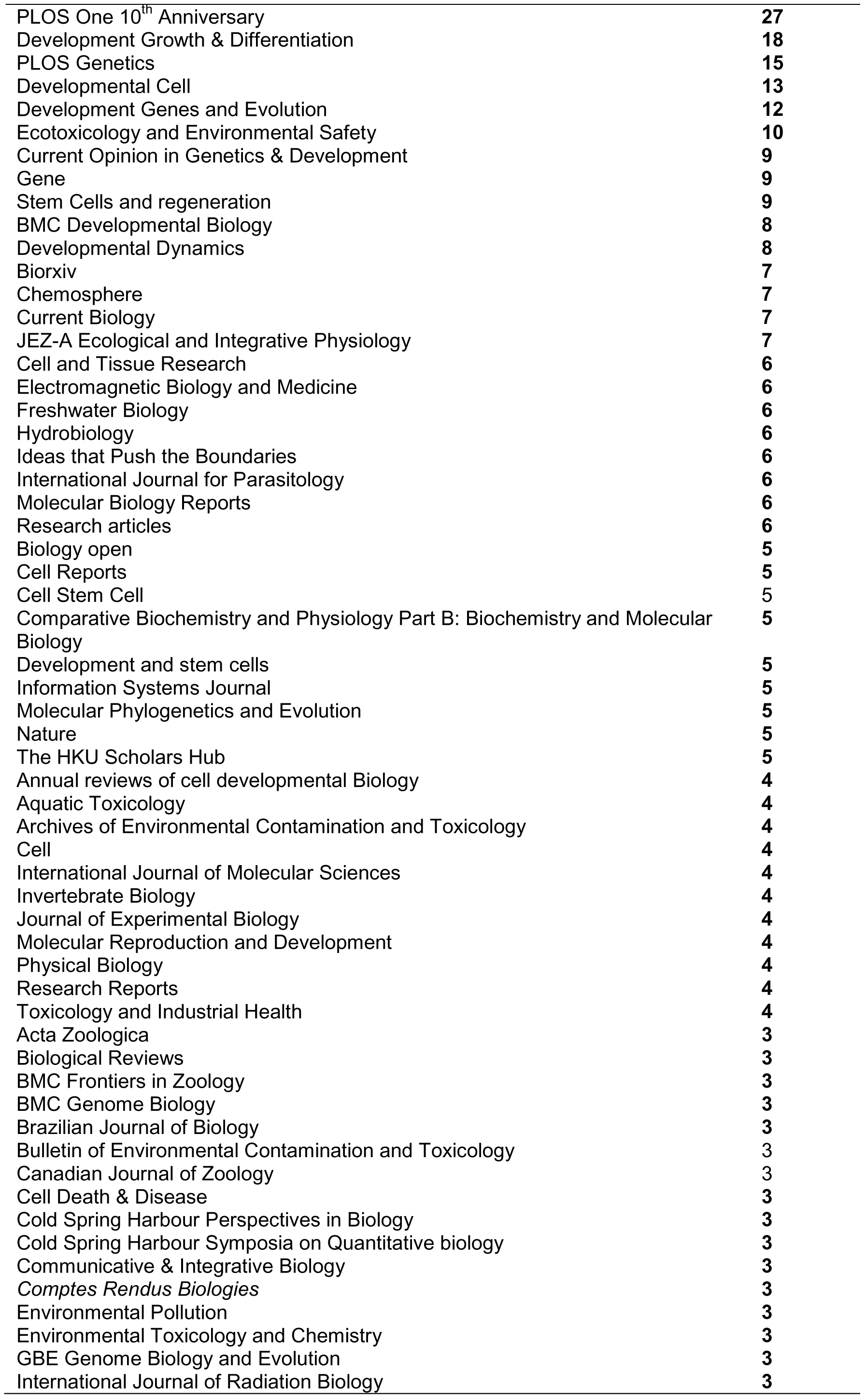
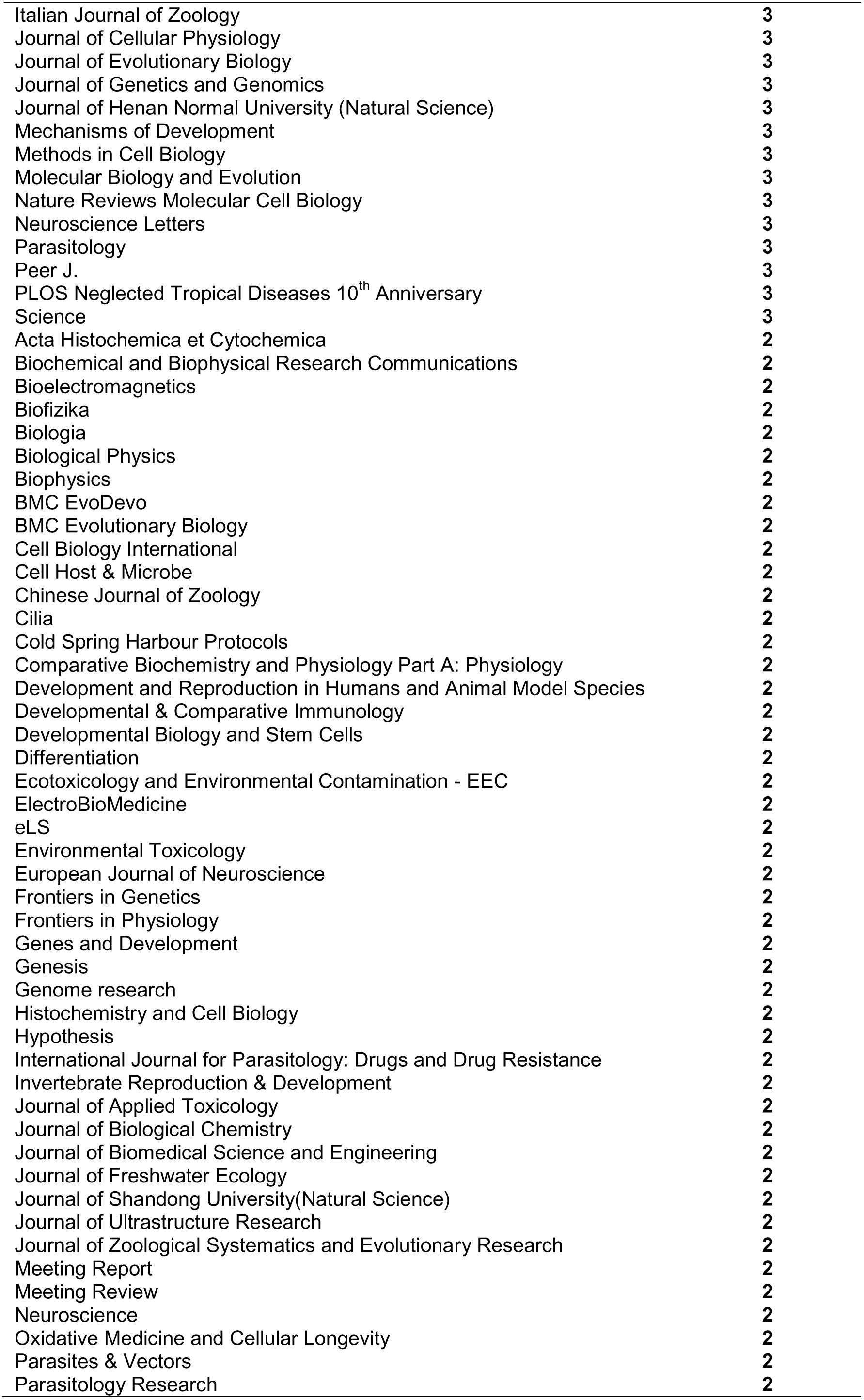
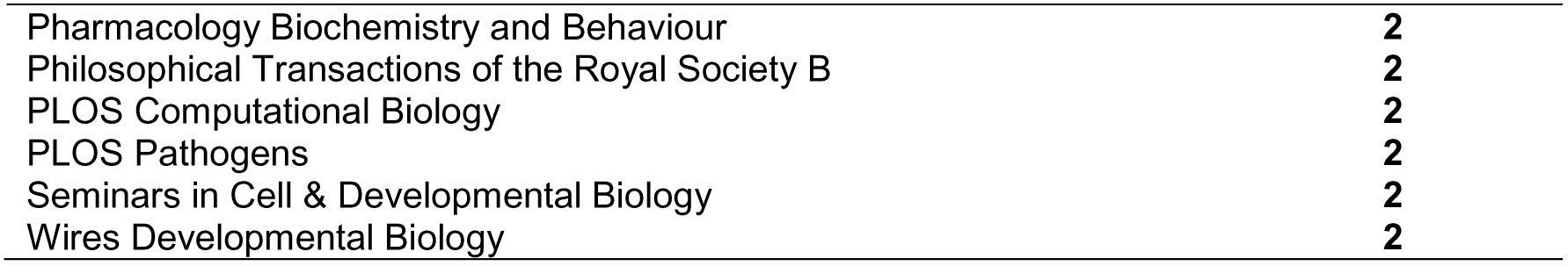
Distribution of papers by journals

|  |  |
| --- | --- |
| Developmental Biology | **30** |
| International Journal of Developmental Biology | **29** |
| PLOS One 10<sup>th</sup> Anniversary | 27 |
| Development Growth & Differentiation | 18 |
| PLOS Genetics | 15 |
| Developmental Cell | 13 |
| Development Genes and Evolution | 12 |
| Ecotoxicology and Environmental Safety | 10 |
| Current Opinion in Genetics & Development | 9 |
| Gene | 9 |
| Stem Cells and regeneration | 9 |
| BMC Developmental Biology | 8 |
| Developmental Dynamics | 8 |
| Biorxiv | 7 |
| Chemosphere | 7 |
| Current Biology | 7 |
| JEZ-A Ecological and Integrative Physiology | 7 |
| Cell and Tissue Research | 6 |
| Electromagnetic Biology and Medicine | 6 |
| Freshwater Biology | 6 |
| Hydrobiology | 6 |
| Ideas that Push the Boundaries | 6 |
| International Journal for Parasitology | 6 |
| Molecular Biology Reports | 6 |
| Research articles | 6 |
| Biology open | 5 |
| Cell Reports | 5 |
| Cell Stem Cell | 5 |
| Comparative Biochemistry and Physiology Part B: Biochemistry and Molecular Biology | 5 |
| Development and stem cells | 5 |
| Information Systems Journal | 5 |
| Molecular Phylogenetics and Evolution | 5 |
| Nature | 5 |
| The HKU Scholars Hub | 5 |
| Annual reviews of cell developmental Biology | 4 |
| Aquatic Toxicology | 4 |
| Archives of Environmental Contamination and Toxicology | 4 |
| Cell | 4 |
| International Journal of Molecular Sciences | 4 |
| Invertebrate Biology | 4 |
| Journal of Experimental Biology | 4 |
| Molecular Reproduction and Development | 4 |
| Physical Biology | 4 |
| Research Reports | 4 |
| Toxicology and Industrial Health | 4 |
| Acta Zoologica | 3 |
| Biological Reviews | 3 |
| BMC Frontiers in Zoology | 3 |
| BMC Genome Biology | 3 |
| Brazilian Journal of Biology | 3 |
| Bulletin of Environmental Contamination and Toxicology | 3 |
| Canadian Journal of Zoology | 3 |
| Cell Death & Disease | 3 |
| Cold Spring Harbour Perspectives in Biology | 3 |
| Cold Spring Harbour Symposia on Quantitative biology | 3 |
| Communicative & Integrative Biology | 3 |
| *Comptes Rendus Biologies* | 3 |
| Environmental Pollution | 3 |
| Environmental Toxicology and Chemistry | 3 |
| GBE Genome Biology and Evolution | 3 |
| International Journal of Radiation Biology | 3 |
| Italian Journal of Zoology | 3 |
| Journal of Cellular Physiology | 3 |
| Journal of Evolutionary Biology | 3 |
| Journal of Genetics and Genomics | 3 |
| Journal of Henan Normal University (Natural Science) | 3 |
| Mechanisms of Development | 3 |
| Methods in Cell Biology | 3 |
| Molecular Biology and Evolution | 3 |
| Nature Reviews Molecular Cell Biology | 3 |
| Neuroscience Letters | 3 |
| Parasitology | 3 |
| Peer J. | 3 |
| PLOS Neglected Tropical Diseases 10<sup>th</sup> Anniversary | 3 |
| Science | 3 |
| Acta Histochemica et Cytochemica | 2 |
| Biochemical and Biophysical Research Communications | 2 |
| Bioelectromagnetics | 2 |
| Biofizika | 2 |
| Biologia | 2 |
| Biological Physics | 2 |
| Biophysics | 2 |
| BMC EvoDevo | 2 |
| BMC Evolutionary Biology | 2 |
| Cell Biology International | 2 |
| Cell Host & Microbe | 2 |
| Chinese Journal of Zoology | 2 |
| Cilia | 2 |
| Cold Spring Harbour Protocols | 2 |
| Comparative Biochemistry and Physiology Part A: Physiology | 2 |
| Development and Reproduction in Humans and Animal Model Species | 2 |
| Developmental & Comparative Immunology | 2 |
| Developmental Biology and Stem Cells | 2 |
| Differentiation | 2 |
| Ecotoxicology and Environmental Contamination - EEC | 2 |
| ElectroBioMedicine | 2 |
| eLS | 2 |
| Environmental Toxicology | 2 |
| European Journal of Neuroscience | 2 |
| Frontiers in Genetics | 2 |
| Frontiers in Physiology | 2 |
| Genes and Development | 2 |
| Genesis | 2 |
| Genome research | 2 |
| Histochemistry and Cell Biology | 2 |
| Hypothesis | 2 |
| International Journal for Parasitology: Drugs and Drug Resistance | 2 |
| Invertebrate Reproduction & Development | 2 |
| Journal of Applied Toxicology | 2 |
| Journal of Biological Chemistry | 2 |
| Journal of Biomedical Science and Engineering | 2 |
| Journal of Freshwater Ecology | 2 |
| Journal of Shandong University(Natural Science) | 2 |
| Journal of Ultrastructure Research | 2 |
| Journal of Zoological Systematics and Evolutionary Research | 2 |
| Meeting Report | 2 |
| Meeting Review | 2 |
| Neuroscience | 2 |
| Oxidative Medicine and Cellular Longevity | 2 |
| Parasites & Vectors | 2 |
| Parasitology Research | 2 |
| Pharmacology Biochemistry and Behaviour | 2 |
| Philosophical Transactions of the Royal Society B | 2 |
| PLOS Computational Biology | 2 |
| PLOS Pathogens | 2 |
| Seminars in Cell & Developmental Biology | 2 |
| Wires Developmental Biology | 2 |

### Most cited papers, authors and countries

It was listed the twenty most cited papers (Table IV) according to the number of cites (over 100 cites) and the correspondent keywords. If the distribution by topics is taken into account, then Stem cell biology and regeneration is the most published topic with eight papers to twenty, followed by molecular and cellular analysis and genetics, both with five papers, and physiological evaluations, morphology and magnetic field, all of them with one paper published.

**Table IV.** Most cited papers and references.

| No. of cites | Authors | Years | Title of the papers | Journal details | Keywords |
| --- | --- | --- | --- | --- | --- |
| 525 | Alvarado, A. S., & Newmark, P. A. | 1999 | Double-stranded RNA specifically disrupts gene expression during planarian regeneration. | *Proceedings of the National Academy of Sciences*, 96(9), 5049-5054. (Alvarado, 1999) | *Planarian* |
| 502 | Reddien, P. W., & Alvarado, A. S. | 2004 | Fundamentals of planarian regeneration. | *Annu. Rev. Cell Dev. Biol.*, 20, 725-757. (Reddien, 2004) | *Planarian* |
| 422 | Newmark, P. A., & Alvarado, A. S. | 2002 | Not your father's planarian: a classic model enters the era of functional genomics. | *Nature Reviews Genetics*, 3(3), 210-219. (Newmark, 2002) | *Planarian* |
| 390 | Reddien, P. W., Oviedo, N. J., Jennings, J. R., Jenkin, J. C., & Alvarado, A. S. | 2005 | SMEDWI-2 is a PIWI-like protein that regulates planarian stem cells. | *Science*, 310 (5752), 1327-1330. (Reddien, 2005) | *Planarian* |
| 345 | Reddien, P. W., Bermange, A. L., Murfitt, K. J., Jennings, J. R., & Alvarado, A. S. | 2005 | Identification of genes needed for regeneration, stem cell function, and tissue homeostasis by systematic gene perturbation in Planaria. | *Developmental cell*, 8(5), 635-649 (Reddien et al., 2005) | *Planaria* |
| 339 | Newmark, P. A., & Alvarado, A. S. | 2000 | Bromodeoxyuridine specifically labels the regenerative stem cells of planarians. | *Developmental biology*, 220(2), 142-153. (Newmark, 2000) | *Planarians* |
| 314 | Wagner, D. E., Wang, I. E., & Reddien, P. W. | 2011 | Clonogenic neoblasts are pluripotent adult stem cells that underlie planarian regeneration. | *Science*, 332(6031), 811-816. (Wagner, 2011) | *Planarian* |
| 283 | Gurley, K. A., Rink, J. C., & Alvarado, A. S. | 2008 | β-catenin defines head versus tail identity during planarian regeneration and homeostasis. | *Science*, 319 (5861), 323-327. (Gurley, 2008) | *Planarians* |
| 267 | Robb, SMC, Ross, E, Alvarado, AS | 2007 | SmedGD: the *Schmidtea mediterranea* genome database | Nucleic acids research 36 (Suppl.-1, D599-606) (Robb, 2007) | *Schmidtea* |
| 244 | Petersen, C. P., & Reddien, P. W. | 2008 | Smed-βcatenin-1 is required for anteroposterior blastema polarity in planarian regeneration. | *Science*, 319(5861), 327-330. (Petersen, 2008) | |
| 244 | Eisenhoffer, G. T., Kang, H., & Alvarado, A. S. | 2008 | Molecular analysis of stem cells and their descendants during cell turnover and regeneration in the planarian *Schmidtea mediterranea*. | *Cell stem cell*, 3(3), 327-339. (Eisenhoffer, 2008) | *Planarian* |
| 231 | Morgan, T. H. | 1898 | Experimental studies of the regeneration of *Planaria maculata*. | *Development genes and evolution* 7(2), 364-397. (Morgan, 1898) | *Planaria* |
| 192 | Meyer, F., Meyer, H., & Bueding, E. | 1970 | Lipid metabolism in the parasitic and free-living flatworms, *Schistosoma mansoni* and *Dugesia dorotocephala*. | *Biochimica et Biophysica Acta (BBA)-Lipids and Lipid Metabolism*, 210(2), 257-266. (Meyer, 1970) | *Dugesia* |
| 192 | Umesono, Y., Watanabe, K., & Agata, K. | 1997 | A planarian orthopedia homolog is specifically expressed in the branch region of both the mature and regenerating brain. | *Development, growth & differentiation*, 39(6), 723-727. (Umesono, 1997) | *Planarian* |
| 170 | Baguña, J., & Romero, R. | 1981 | Quantitative analysis of cell types during growth, degrowth and regeneration in the planarians *Dugesia mediterranea* and *Dugesia tigrina*. | *Hydrobiologia*, 84(1), 181-194. (Baguña, 1981) | *Dugesia* |
| 166 | R Thompson, J McConnell | 1955 | Classical conditioning in the planarian *Dugesia dorotocephala*. | *Journal of comparative and physiological psychology*, 48(1), 65. (Thompson and McConnell, 1955) | *Dugesia* |
| 142 | Rink, J. C. | 2013 | Stem cell systems and regeneration in Planaria. | *Development genes and evolution*, 223(1-2), 67-84. (Rink, 2013) | *Planaria* |
| 123 | FA Brown Jr. | 1962 | Responses of the planarian, *Dugesia*, and the protozoan, *Paramecium*, to very weak horizontal magnetic fields | *The Biological Bulletin*, 123(2), 264-281. (Brown Jr, 1962) | *Dugesia* |
| 108 | Adamidi, C., Wang, Y., Gruen, D., Mastrobuoni, G., You, X., Tolle, D. & Rybak, A. | 2011 | De novo assembly and validation of planaria transcriptome by massive parallel sequencing and shotgun proteomics. | *Genome research*, 21(7), 1193-1200. (Adamidi et al., 2011) | *Planaria* |
| 102 | Curtis, W. C. | 1902 | The life history, the normal fission and the reproductive organs of *Planaria maculata* | Boston Society of Natural History from the Gurdon Saltonstall Fund. Vol. 30, No. 7. (Curtis, 1902) | *Planaria* |

It also analysed the paper distribution by journal. There are16 journals which published the most cited papers. The journals that have more papers published about Planarian research since 1955 to 2017 is Science, with only four papers published, and Development genes and evolution with two papers. Concerning with the distribution *per* year taking into account the three stages previously defined; the 2001-2010 stage had the most relevant papers (8), followed by 1955-2000 (7), and the 2011-2017 (3). There were two papers before 1955 (Table IV).

The most cited paper (and author) was: ¨Double-stranded RNA specifically disrupts gene expression during planarian regeneration¨ (Alvarado & Newmark, 1999) with 525 cites. These two authors and Reddien were the most cited authors in the twenty papers chosen.

Otherwise, it is important to mention that the species reported in these researches were: *Schmidtea mediterranea, Planaria maculate and Dugesia dorotocephala, Schisostoma mansoni*, *D. mediterranea* and *D. tigrina*.

### Distribution of papers by countries

This analysis was based on the country by filiations, it means the country of the institution of main author in each paper (n=788) (Fig. 4).

**Fig. 4.**
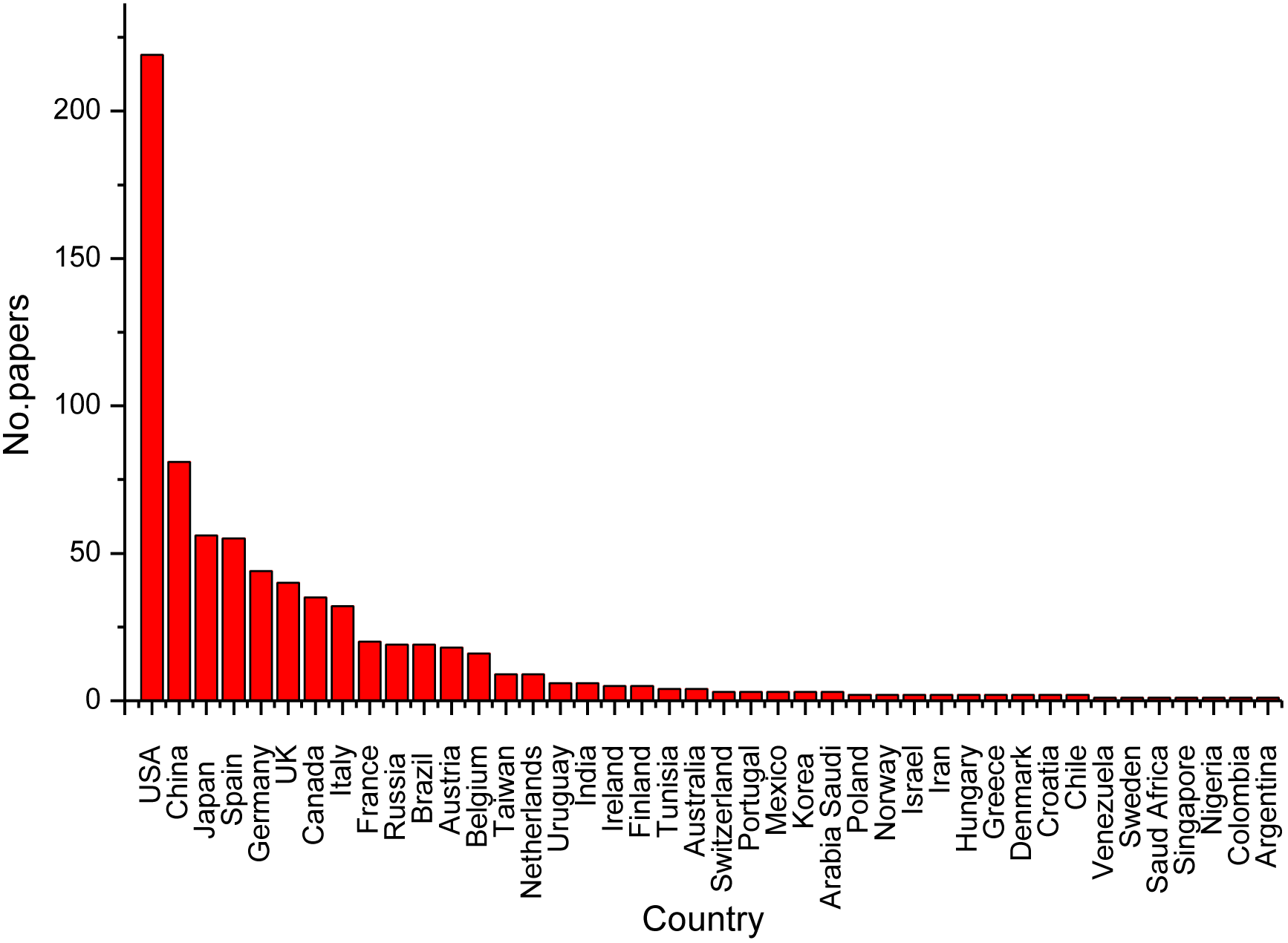
Distribution of papers by countries, n=788.

As a general comment there are a group of eight countries with an important contribution in Planarian research, being USA the most relevant with more than 200 papers, followed by China, Japan and Spain with more than 50 but less than 100, and Germany, UK, Canada and Italy with more than 25 but less than 50. It means that these researches are concentrated in developed countries, mainly.

### Analyses of Planarian research focused on magnetic field

Considering that the real interest which conducted this meta-analysis was to know the state of the art concerning with magnetic field using Planarian as biological model, a number of 57 papers as a specific search result were analysed. It was focused on magnetic field researches, but the majority was related with other biological models, but mention the results on Planarian. The final analysis allows selecting nine papers, published during the last three years. It is an evidence that this topic is becoming a trending with rising interest (Table II).

The most reported species to study the magnetic field effects on Planarian were mainly *Dugesia* spp. which is referred in five papers, while *Girardia* sp. appears in just one paper, both belong to Dugesiidae family.

Concerning with magnetic field types and doses, the electromagnetic field (EMF) was used in seven papers (78%) and static magnetic field (SMF) in two.

Weak electromagnetic field (1 to 5 μT, 38 Hz) was used in three papers; pulsing electromagnetic field (PEMF) or weak pulsing electromagnetic field 5 μT (8, 16 or 72 Hertz) in two papers; extremely low frequency electro-magnetic field (ELF-EMF) (60 Hz and 80 mG, applied twice a day for one hour) in one paper; while other papers deals with the geomagnetic patterned MF 7 Hz range (50 nT or 200 nT); the weak temporally-patterned magnetic field (0,1 to 2 and 0,5 to 5 μT) and the last one about the effect of SMF (16, 160, or 1,600 G).

In order understand the research strategies the main objectives of these nine researches were analysed. These were focus in regeneration (5) and fragmentation processes (1), stress protein expression (2), interaction of pigments (melatonin) with MF (1) and behavioural response or Planarian activity (2); all of these are listed below:

1. to study the regeneration phenomena in a frequency modulated patterned EMF in *D. tigrina* (Tessaro and Persinger, 2013);
2. to increase the regeneration associated with marker activation of the extracellular signal regulated kinases and heat shock protein (Murugan, 2013);
3. to determine the effect of week combined MF tune to the cyclotron resonance condition for calcium ions obtained at different faces of Planarian regeneration (Tiras et al., 2015);
4. to investigate beneficial effects of early rotary none uniform MF of 6 Hz (TMF) exposure on central nervous regeneration in *Girardia sinensis* (Chen et al., 2016)
5. to use regenerating Planaria *D. dorotocephala* as a model to determine whether an intermittent modulated ELF EMF produce elevated levels of the heat shock protein hSP 70 and stimulated intracellular pathways none to be involved in injury and repair (Goodman et al., 2009);
6. to evaluate if the temporally-patterned magnetic fields (0,1-2μT and 0,5-5 μT) induce complete fragmentation (Murugan et al., 2013);
7. to test the synergistic interactions between specific MF (7 Hz) and chemical concentration of melatonin using *Dugesia* sp.(Mulligan et al., 2012);
8. to test the behavioural responses of Planaria to the exposure of a range of concentration of morphine, naxolone or to either of these compounds and a burst-fighting MF 5 μT (Murugan and Persinger, 2014);
9. To evaluate the effect on Planarian activity of maintained water pre-treated with magnetic fields (Gang and Persinger, 2011).

It is important to remark that the 89% (eight of nine) of paper were published since 2012 to 2017, and the other paper in 2009.

As a most important results of the EMF application presented in these papers are the increasing activity of mARN with the application of 60 Hz WMF, while WMF less than 1 μT accelerated the oxidation of cytochrome C *in vitro* and affect the function of Na/K+ ATPase (Murugan, 2013).Other results highlight the multiplicity of enzymatic targets activated at different phases of the regeneration process, 72 h after decapitation, which depend on the exposure duration and the time between decapitation and initiation of the exposure (30 min.) with a week combined MF tune to the cyclotron resonance (Tiras et al., 2015).

It is important to mention too, that at present, there are many data about the weak combined (alternating and constant) magnetic fields (CMF) with an amplitude close to that of the Earth’s magnetic field, capable of inducing changes in metabolism of different living organisms, from human to microorganisms (Liboff, 2013).

Otherwise Planarian (*Dugesia tigrinia*) exposed to a frequency-modulated (“Thomas”), patterned electromagnetic field (EMF) immediately following transection through the pharynx with an exposure from 15 min to 3 h as well as single *versus* repeated exposures, resulted in a better regeneration rates that in those planaria exposed from 45 to 90 min. In addition, the study revealed that exposures greater than 45 min were not significantly different beyond this inflection point, suggesting that this particular pattern of EMF is capable of inducing biochemical pathways associated with cell proliferation, in particular the p38-MAPK and hsp70 pathways (Tessaro and Persinger, 2013).

One of the results is related with changes of Planarian´s volume in specimens maintained in spring water and exposed for two hours to temporally patterned, weak (1 to 5 μT) magnetic field in the dark. It diminished mobility that simulated the effects of morphine and enhanced this effect at concentrations associated with receptor subtypes. A single (5 hr) exposure to this same pattern following several days of exposure to a very complex patterned field in darkness dissolved the planarian associated with an expansion of their volume. Continuous measurement of pH indicated that the slow shift towards alkalinity over 12 hours of exposure was associated with enhanced transient pH shifts of 0.02 units with typical durations between 20 and 40 ms. These results indicate that the appropriately patterned and amplitude of magnetic field that affects water directly, could mediate some of the powerful effects displayed by biological aquatic systems (Murugan, 2013).

A tandem sequence composed of weak temporally-patterned magnetic fields was discovered that produced 100% dissolution of planarian in their home environment. The expansion (by visual estimation ∼twice normal volume) of the planarian following the first field pattern followed by size reduction (∼1/2 of normal volume) and death upon activation of the second pattern was observed. It suggest effects on contractile proteins and alterations in the cell membrane’s permeability to water due to the MF effect (Murugan et al., 2013). In our laboratory dissolution of marine microturbelarian was observed in a tandem sequence of experiments with SMF (unpublished data).

To test the hypothesis that there are some results which explain that exposing water to a static magnetic field affects its properties which influence living systems, planarian subsequent to dissection were maintained in spring water that had been previously exposed for one day to one of three (16, 160, or 1,600 G) intensity static magnetic fields or to a reference condition. Although there was no significant difference in regeneration rates over the subsequent seven-day period, there was a statistically significant nonlinear effect for planarian mobility and diffusion rates. Both mobility rates and diffusion velocity of a liquid within the water that had been exposed to the 16 G field was about twice that for water exposed to the other intensities. These results imply that nonlinear biophysical effects may emerge under specific conditions of intensity ranges for particular volumes of water (Gang and Persinger, 2011).

Research concerning with the exposures to a modulated sinusoidal ELF-EMF delivered by a Helmholtz configuration at a frequency of 60 Hz and 80 mG twice a day for one hour. This is accompanied by an increase in hsp70 protein levels, activation of specific kinases and up regulation of transcription factors that are generally associated with repair processes. During the initial 3-days post-surgery caused a significant increase in regeneration for both heads and tails, but especially tails. The first appearance of eyes occurred at day seven post-transection in tail portions exposed. In the sham control tail samples, the initial appearance of eyes occurred 48 h later. Concurrently, ELF-EMF-exposed heads and tails exhibited an elevation in the level of hsp70 protein, an activation of an ERK cascade, and an increase in SRF-SRE binding (Goodman et al., 2009).

A significant knowledge gap persists between molecular genetics components identified as necessary to produce a particular organism shape, regeneration process and understanding how and why a particular organism with a simple or complex shape has the ability to generated new tissues to regenerate the initial shape in the correct size, shape and orientation; otherwise there are a deep gap in the understanding of the effect of magnetic field in regeneration processes, genetic implication, proteomics, and action mechanisms, and Planarian use to be an adequate biological model to develop these studies. More researches are needed to clarify all of these questions and other which arise during the following findings, but definitively the magnetic field effect on Planarian is a wide field of action with many challenges and mysteries to highlight.

## Acknowledgements

We would like to thank to the professors Tom Artois and Yander Diez García, for inspire us to work with Planarian and marine microturbelarians, providing the first specimens and to introduce us in the main aspects of these organisms.

## Author contributions

All authors performed the meta-analysis and wrote the manuscript.

